# Single-cell characterization of step-wise acquisition of carboplatin resistance in ovarian cancer

**DOI:** 10.1101/231548

**Authors:** Alexander T. Wenzel, Devora Champa, Hrishi Venkatesh, Si Sun, Cheng-Yu Tsai, Jill P. Mesirov, Jack D. Bui, Stephen B. Howell, Olivier Harismendy

## Abstract

Acquired resistance to carboplatin is a major obstacle to the cure of ovarian cancer, but its molecular underpinnings are still poorly understood and often inconsistent between in vitro modeling studies. Using sequential treatment cycles, multiple clones derived from a single ovarian cancer cell reached similar levels of resistance. The resistant clones showed significant transcriptional heterogeneity, with shared repression of cell cycle processes and induction of IFNα response signaling, and subsequent pharmacological inhibition of the JAK/STAT pathway led to a general increase in carboplatin sensitivity. Gene-expression based virtual synchronization of 26,772 single cells from 2 treatment steps and 4 resistant clones was used to evaluate the activity of Hallmark gene sets in proliferative (P) and quiescent (Q) phases. Two behaviors were associated with resistance: 1) broad repression in the P phase observed in all clones in early resistant steps and 2) prevalent induction in Q phase observed in the late treatment step of one clone. Furthermore, the induction of IFNα response in P phase or Wnt-signaling in Q phase were observed in distinct resistant clones. These observations suggest a model of resistance hysteresis, where functional alterations of the P and Q phase states affect the dynamics of the successive transitions between drug exposure and recovery, and prompts for a precise monitoring of single-cell states to develop more effective schedules for, or combination of, chemotherapy treatments.

## Introduction

Patients diagnosed with high grade serous ovarian cancer are generally treated initially with either cisplatin or carboplatin (CBDCA) in combination with paclitaxel. While 65-75% of patients respond to the primary treatment ^1^, resistance emerges frequently during therapy and this is a major obstacle to cure ^2^. Unlike targeted agents where high-levels of resistance are common, repeated treatment of sensitive cells with clinically relevant levels of exposure to cisplatin or CBDCA produces only low-level resistance, typically in the range of 1.5-3-fold, a level sufficient to account for clinical failure of treatment *in vivo* ^3^.

The mechanisms underlying acquired resistance to platinum-containing drugs have been the subject of intense study ever since their discovery. Acquired resistance has been attributed to changes in many types of cellular functions including import and export of the drug, enhanced detoxification and DNA adduct repair, inactivation of the mismatch repair checkpoint and repression of apoptotic signaling ^4^. Findings from single genes or transcriptome-wide studies of bulk cell populations can usually be validated through overexpression or knock-out, but these studies have failed to disclose any actionable gene or set of genes that are consistently altered across different cell types or experiments and that would point toward the need for widely useful approaches for preventing or overcoming the development of resistance in patients.

Apart from rare instances of *BRCA1/2* mutation reversion^5^, the acquisition of CBDCA resistance is believed to be epigenetically mediated ^6^. Recent advances in the study of resistance to kinase inhibitors has revealed the existence of “persister” cells in lung cancer cell lines that are present at low prevalence and can resist treatment through epigenetic mediated mechanisms ^7^. Previously, single cell tracing had shown that, within a cell population, the immediate response to genotoxic treatment can vary extensively from cell to cell giving rise to considerable heterogeneity within the surviving population ^8,9^. More recently, similar observations were made in vemurafenib-treated melanoma cells where cells in a transient resistant state are present in the population prior to drug exposure and display increased levels of expression of resistance genes ^10^. Importantly, the relevance of these recent models to the acquisition of resistance to platinum-containing drugs has not been established for either *in vitro* or *in vivo* models nor have the concepts been validated in clinical studies.

Here we present a comprehensive phenotypic and molecular characterization of a set of ovarian cancer clones derived from a single cell and selected in parallel for acquired resistance to CBDCA. We show that the resistance is unlikely to be due to genetic mutations, copy number changes or differences in CBDCA uptake. Resistance was associated with significant changes in proliferation rate and the capacity to form colonies and organoids, and there is substantial heterogeneity between clones. Transcriptome profiling showed a common association of CBDCA resistance with slow proliferation and high interferon signaling but also demonstrated marked heterogeneity between resistant clones. Importantly, single-cell transcriptome analysis allowed us to characterize the resistant states at single-cell resolution, eliminating the confounding effect of cell cycle variation and revealing functional changes specific to proliferative and quiescent phases.

## Results

While resistance acquisition can be well recapitulated *in vitro* through repeated drug exposure, variation between cell lines, methodologies and the lack of selection replicates compromise the identification of widely shared molecular changes associated with resistance to platinum-containing drugs. We chose an experimental design that would allow us to determine whether genetically identical clones undergo the same molecular changes during acquisition of resistance. Specifically, we isolated a single cell from a non-clonal population of human ovarian cancer CAOV3 cells, and grew them to a small population (the parental clone) from which 12 clones were isolated (Figure 1A). Four of these clonal populations were grown continuously in the absence of CBDCA (“S clones”: S01-S04). The remaining 8 (“R clones”: R06, R07, R14-R19) were each individually subjected to 4 cycles of exposure to CBDCA at which point they tolerated 5 µM drug (subsequently referred to as step 5) and averaged 1.7-fold resistance relative to the parental clone (Figure 1B). Four of these 8 clones were then treated with additional cycles of CBDCA at gradually increasing concentrations until they tolerated 15 µM CBDCA (referred to as step 15) and averaged 7.8-fold resistance relative to the parental clone. Subsequent passages of the resistant clonal populations in the absence of CBDCA for 63 doublings did not result in loss of resistance (Figure S1), indicating that the phenotype was stable within the number of passages used in the study.

**Figure 1:**
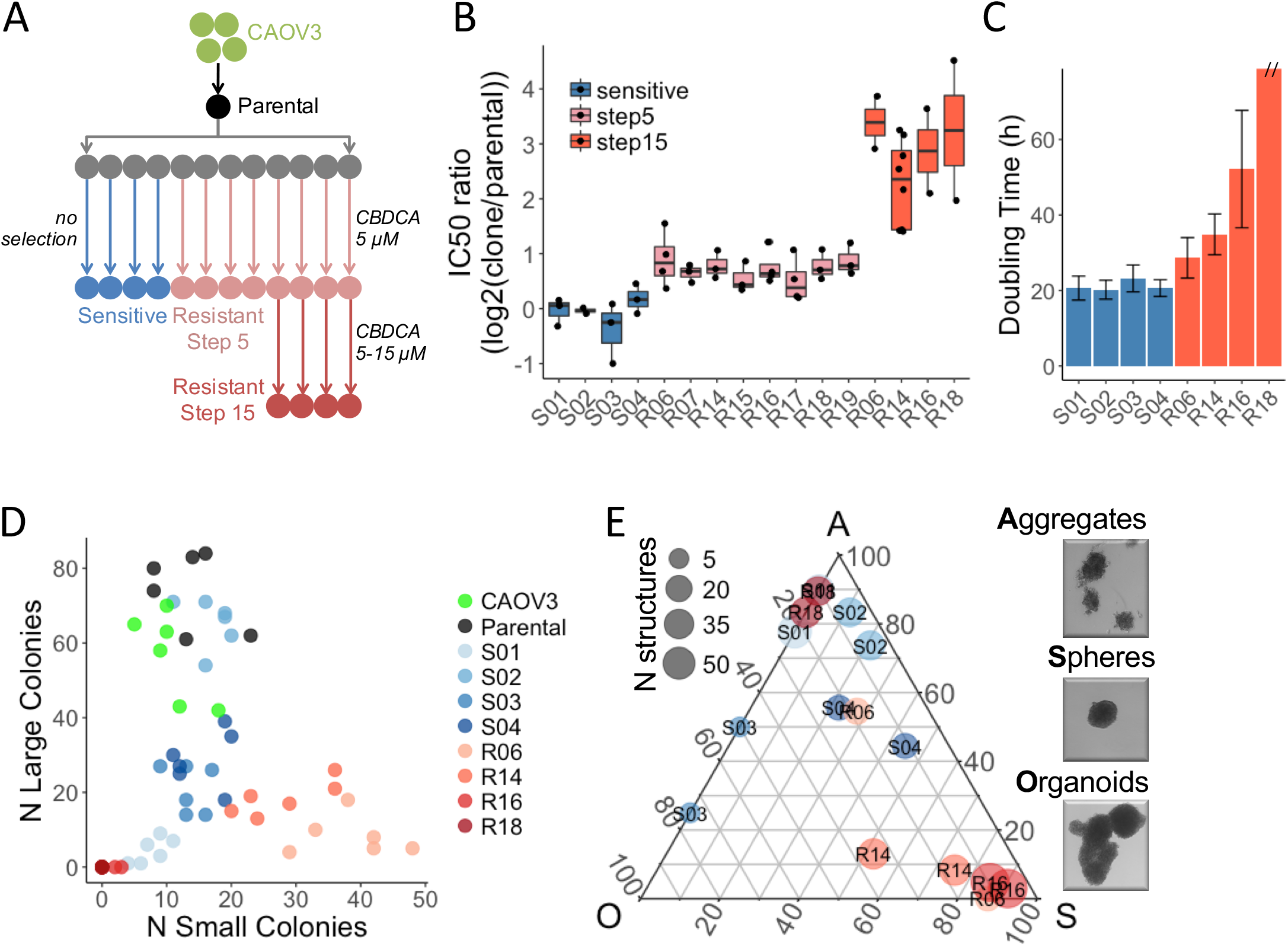
Phenotypic characterization of the resistant clones. **(A)** Schematic representation of the workflow to generate CBDCA resistant clones from CAOV3. **(B)** Changes in IC50 of S clones (unselected) or R clones (8 at step 5 and 4 at step 15). Each IC50 is calculated from dose-response curves of 6 replicates and experiments repeated twice or more (dots). **(C)** Doubling time measured over a 48 h time course – y axis cut for R18 (>100 h). **(D)** Counting of colonies formed in a period of 9 days after seeding 200 cells per well. Experimental replicates (N=6) are shown. **(E)** Fraction of organoids (O), spheres (S) and cell aggregates (A) observed after 14 days growth in low adherence 3D culture model. For each sample (N=8) and replicates (N=2), the total number (point size) and relative abundance (Gibbs triangle coordinates) of each type of structure are indicated.

### Phenotypic and molecular characterization of isogenic resistant clones

We next compared the phenotypic and molecular characteristics of 4 S and 4 step 15 R clones to identify features associated with acquired resistance. As shown in Figure 1C-E, the growth of the R clones was slower, they formed fewer large colonies in 2D culture, and in low-attachment plates they formed a higher proportion of small spheres. Importantly, the reduced proliferation can partly explain the resistant phenotype (Figure S2) as previously proposed^11,12^. However, given the large differences in proliferation between R clones despite their similar level of resistance, it is likely that other processes contribute to the reduced cytotoxicity. Cells from all clones had similar distribution of CBDCA content after 1 h exposure (Figure S3), suggesting that unlike other models^13^, reduced drug uptake was not a major contributor to resistance in the studied clones. Exome sequencing of 4 S clones and 8 R clones (step 5) was used to identify copy number alterations (Table S1). Two clones (S01 and R06) were affected by copy number gains (4 and 10 Mbp) and 7 clones (S01-03, R06, R16, R18, R19) had copy number losses (0.2-12 M bp). When copy number changes occurred, they were small in magnitude (less than 1.5-fold) and none of them affected multiple R clones. We also identified a median of 39 coding mutations per clone affecting a total of 74 genes. Neither total mutation burden nor gene-specific mutational burden was significantly associated with the resistant phenotype, albeit with limited statistical power (Table S2). Interestingly, *CAAP1* T103P, was identified in all 8 R clones and 1 S clone, and while the mutation is predicted to be deleterious (CADD score =22^14^), a role for *CAAP1* in apoptosis signaling has not been validated ^15,16^. Thus, these clonal populations derived from a single cell were genetically homogeneous and the small numbers of genetic changes observed were unlikely to be causally associated with the resistant phenotype.

In order to identify molecular processes associated with resistance, we measured the expression level of all genes in the 4 S and the 4 R clones at step 15 using RNA-seq. We identified 186 genes that were differentially expressed between S and R clones (Figure 2A). An enrichment analysis of Hallmark ^17^ and Reactome ^18^ gene sets from MSigDB^19^, revealed that resistance was associated with a global repression of proliferation and translation and the activation of genes involved with interferon and *KRAS* signaling, and epithelial-to-mesenchymal transition (EMT) (Figure S4). All of these are processes previously reported to be involved in chemo-resistance or response to genotoxic injury^20–22^. An unsupervised analysis revealed that, while the S clones had similar transcriptional profiles, the profiles of the R clones were highly heterogeneous (Figure 2B). Processes related to cellular proliferation (E2F targets) were repressed in all R clones, and interferon and *KRAS* signaling were induced in all R clones (Figure 2C). In contrast, a clone-specific analysis revealed that cell cycle and nucleotide excision repair were induced at higher level in R06, EMT in R14, oxidative phosphorylation in R16. These processes are not mutually exclusive and were dysregulated to different degrees in the various resistant clones. Importantly, the regulation of these resistance-associated processes is not specific to CAOV3. The analysis of gene expression profiles associated with acquired cisplatin resistance in 8 ovarian cancer cell lines^23^, confirms the greater heterogeneity of resistant clones compared to untreated cells and shows that genes involved in IFNα or KRAS signaling, or EMT are frequently activated and those related to cell cycle and proliferation are repressed at equivalent time point and treatment regimen (cycle 6, schedule C, Figure S5). Interestingly, we observed that ruxolitinib, a strong inhibitor of IFNα signaling via the JAK/STAT pathway, could re-sensitize the CAOV3 cells to CBDCA (Figure 2D) suggesting that the JAK/STAT pathway activation is required for the resistance in these cells. Similarly, inhibition of *KRAS* signaling via silencing of MEK or inhibition of ERK1/2 has been shown to increase platinum sensitivity in other ovarian cancer cell lines^24–26^. Beyond common mechanisms of resistance highlighted or confirmed by this analysis, our observations suggest important differences between clones prompting a more detailed investigation of the dynamic changes in the processes mediating treatment resistance.

**Figure 2:**
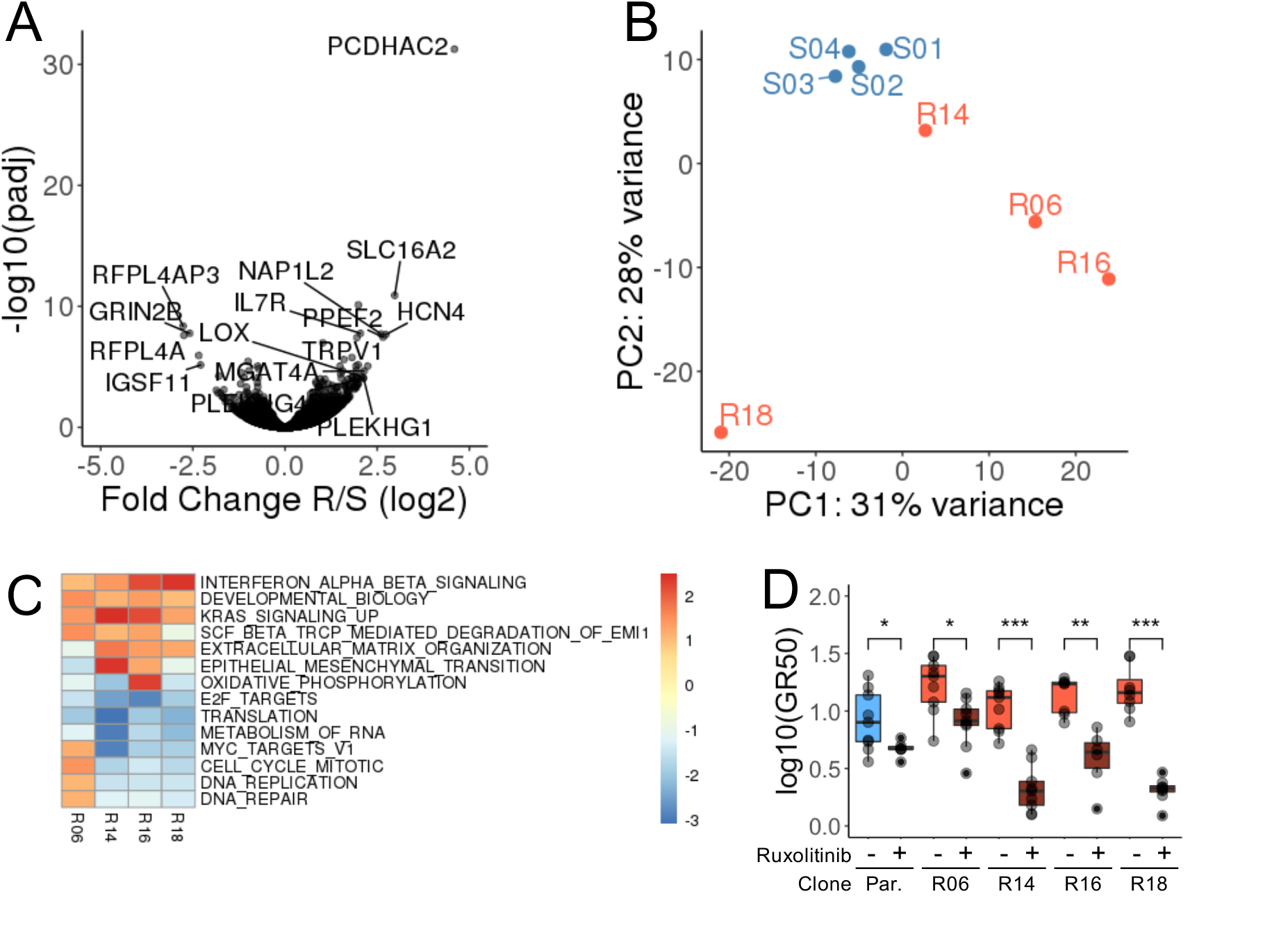
Expression profiling of the derived clones. **(A)** Volcano plot indicating the fold change (y axis) and significance (x axis) of the genes differentially expressed between S and R clones. **(B)** First two principal components derived from the expression profiles of each clone. **(C)** Most significantly up or down-regulated gene sets (Hallmark and Reactome from MSigDB) in individual R clones compared to all S clones. Significant gene sets (q.value<0.005) enriched (score>1.5) or depleted (score <-2) in at least one clone are reported. Color gradient indicates enrichment score. **(D)** Treatment with 5 µM ruxolitinib (Rux) significantly decreases the growth-rate corrected half maximal inhibitory concentration (GR50) in both Parental and R clones. The results of 3 dose-response experiments, 3 replicates per experiment are presented. Significance was measured using Wilcoxon Test (*<0.05, **<0.01, ***<0.001).

### Characterization of chemo-resistance at single cell resolution

Suspecting that phenotypic heterogeneity within the cell population may be underlying the differences in the acquisition of drug resistance, we measured the expression level of individual genes in 26,772 single cells from the original CAOV3 cell population, the parental clone derived from a single cell in this population, two S clones and two time points for each of the R clones (step 5 and step 15). The cells were classified according to their cell cycle signatures and, in agreement with the slower proliferation of the resistant clones, step 5 resistant cells were more likely to be quiescent than the untreated cells (61% vs 21% in G0-Figure 3A), while step 15 cells resumed proliferation (∼30% in G0) with the exception of R16 (57% in G0).

**Figure 3:**
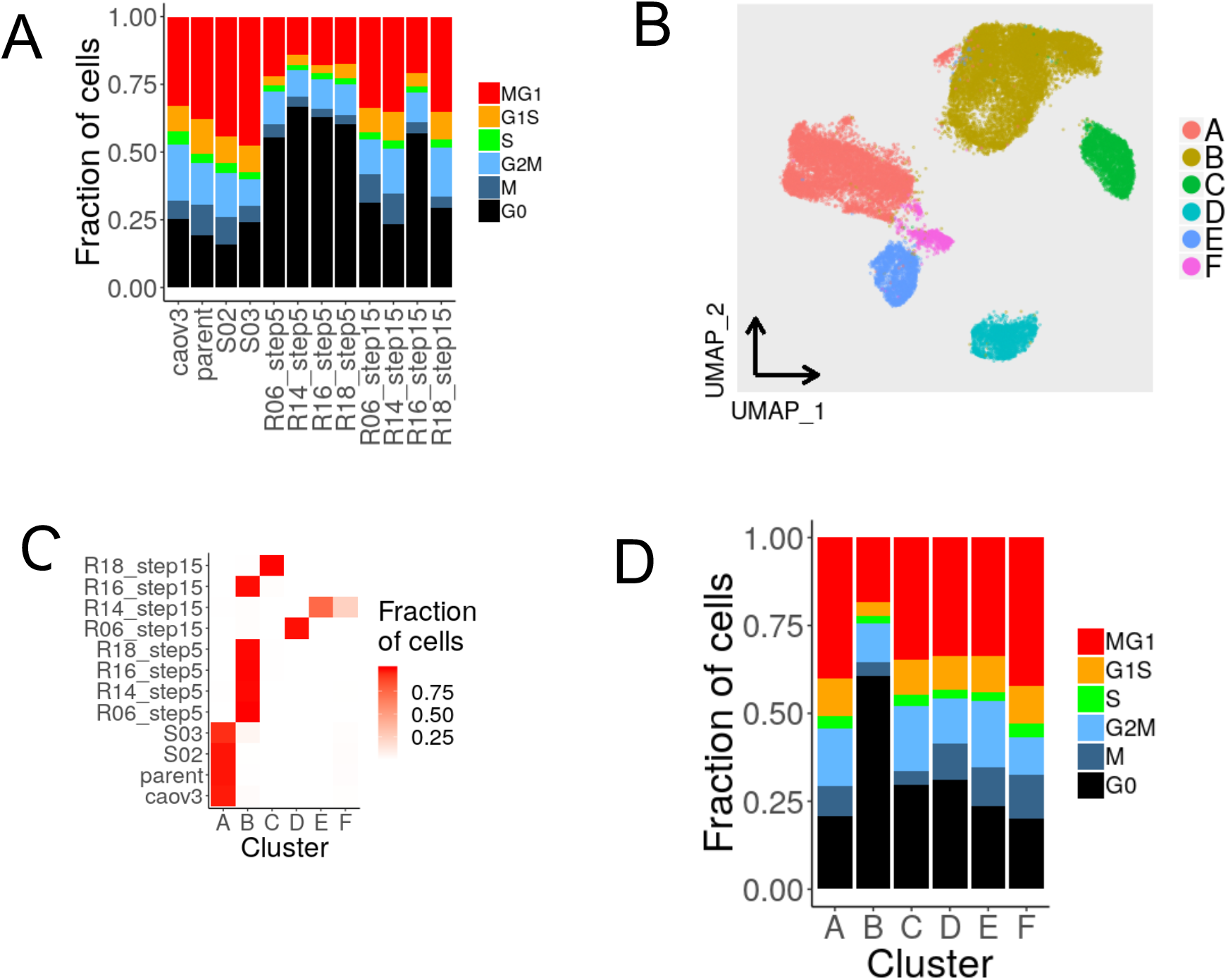
Evolution of expression states in all clones. **(A)**. Distribution of cells in three phases of the cell cycles estimated from the expression signatures. **(B)** Uniform Manifold Approximation and Projection (UMAP) of cells from the aggregated analysis based on the first 2 principal components. Cells are colored according to the Louvain clusters. **(C)** Distribution of the cells from each clone and treatment group across the 6 clusters. **(D)** Distribution of cells in three phases of the cell cycles estimated from the expression signatures.

The expression profiles were used to group all cells into 6 expression clusters, A through F (Figure 3B). Cells from step 15 R clones were distributed across 5 different clusters, whereas cells from step 5 R clones clustered together, indicating that additional treatment cycles increase transcriptional heterogeneity (Figure 3C). Interestingly, cells from R14 step 15 cells likely split into two sub-populations which could not be reliably distinguished by chromosomal copy number analysis (Figure S6A) suggesting their differences in expression are not genetically driven. Similarly, step 15 clones display increase aneuploidy compared to their step 5 clones, suggesting that genetic changes may contribute to their increased heterogeneity (Figure S6B-C). The single-cell analysis further revealed that a small fraction (230/7814, 2.9%) of untreated cells were not in cluster A and were primarily in cluster B (133/230, 57%). The converse was not true with fewer than 0.2 % of the treated cells found in cluster A. This observation suggests that untreated cells exist in a pre-resistant state prior to exposure to the drug. Cluster B had the largest fraction of cells in G0 indicating that quiescence characterizes both pre-resistant cells, step 5 and R16 step 15 cells (Figure 3D). However, differences in cell-cycle alone is unlikely to explain the variation in cellular states as cells in all phases of the cell cycle exist in all clusters. This observation prompted us to more carefully account for cell cycle differences before characterizing the functional changes associated with the different resistant states.

### Variation of expression-based activity of biological processes along the cell-cycle

The contribution of cell cycle to single-cell gene expression can be mathematically subtracted to study the source of the residual variation and its association with the resistant phenotype. However, this correction method assumes the independence of gene expression measurements. Nonetheless, gene expression is tightly coordinated and gene-based cell-cycle correction may mask important intrinsic differences as cells progress through the cell cycle. Thus, to make the correction, we chose instead to use the expression of 603 genes associated with cell cycle to order the cells along a linear pseudotime (pt) trajectory from mitosis (G2M, M; pt < 0.20) to growth and replication (MG1, G1S, S; 0.20 ≤ pt < 0.69) and quiescence (G0; pt ≥ 0.69) (Figure 4A-B). Consistent with the cell cycle phase analysis, the distribution of cells along the trajectory varied between clusters with the majority of cells in cluster A (referred to as A cells) in proliferation (8% with pt≥0.69), B and C cells mostly in quiescence (79% and 74% pt≥0.69 respectively), while D, E and F cells, corresponded to step 15 treatment of R06 and R14 clones, resuming proliferation (19%, 15% and 15% with pt≥0.69, respectively). The resulting virtual synchronization allowed us to identify cells from different clones that are in similar phases of the cell cycle, which facilitates the interpretation of functional differences associated with resistance.

**Figure 4:**
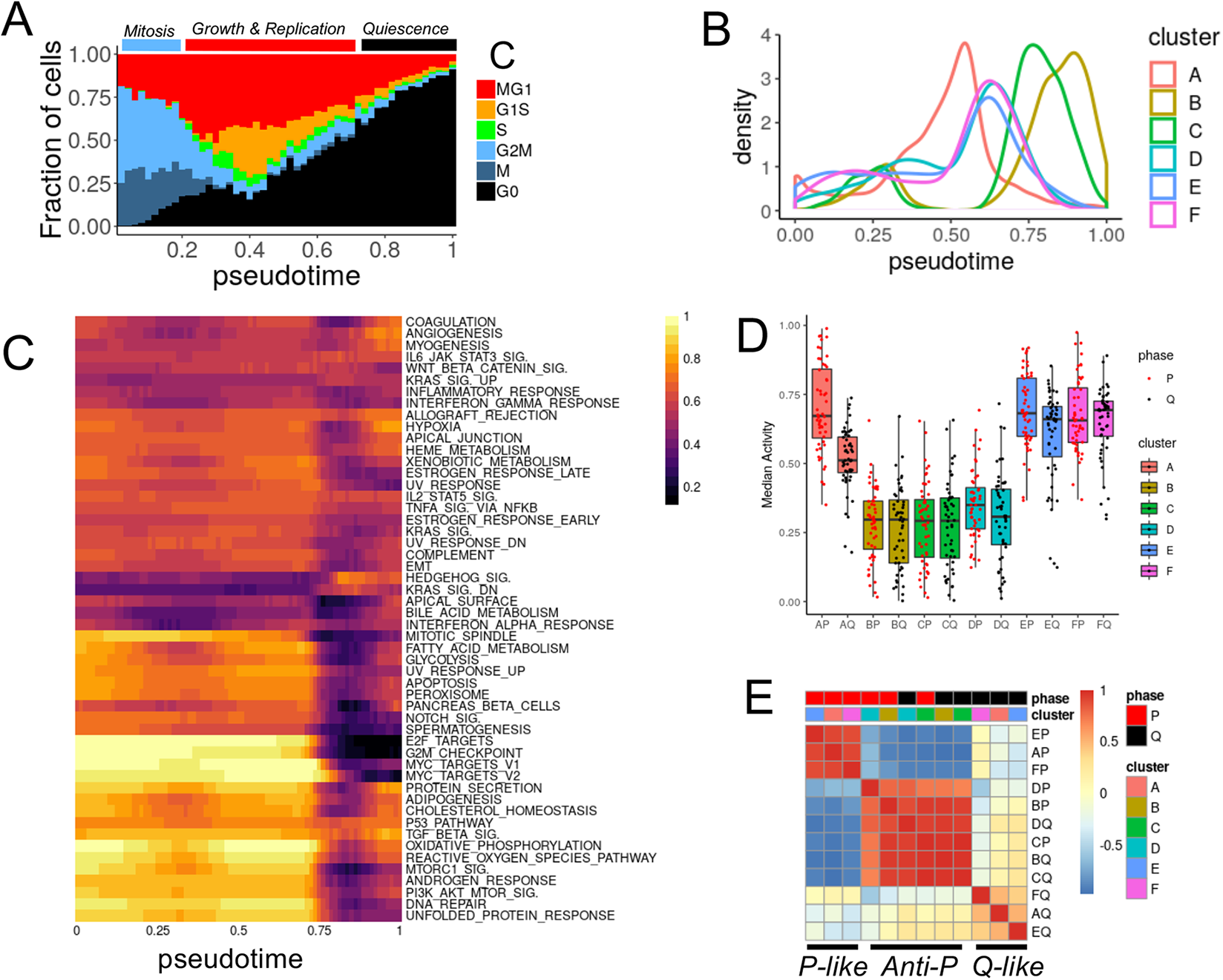
Functional analyses of single-cell resistant states after in silico synchronization. **(A)** The distribution of cells (kernel density – y axis) along the pseudo-time trajectory (x axis) is represented for each expression-based clustering. **(B)** Fraction of cells in each inferred cell cycle phase as a function of the pseudo-time (x axis bins). **(C)** Scaled enrichment score (ES) of the hallmark gene sets (clustered rows) observed in cluster A cells as a function of pseudo-time (columns). **(D)** Median scaled enrichment score of all hallmark gene sets across cells from P (red points) and Q (black points) phases for each of the 5 clusters. **(E)**. Correlation of hallmark gene sets median scaled enrichment score for all clusters and proliferation phases.

The changes in activity, or enrichment score (ES), of the Hallmark gene sets can be measured along the pseudo-time using the smoothed geometric average of the gene expression (see methods – Figure 4C). In untreated cells (Cluster A), the activity varied most strongly between proliferation (referred to as P; pt<0.69) and quiescence (referred to as Q; pt≥0.69). In particular, and as expected from their use in the trajectory inference, processes associated with cell cycle progression had the highest activity in P and shut down in Q (G2M checkpoint, E2F target, MTOR signaling). The activity of other processes were either unchanged along the cell cycle (TGFβ signaling, P53 pathway) or were induced as cells progressed from P to Q (Hedgehog signaling, KRAS_DN). These variations, combined with the strong proliferative differences

between clones, highlight the necessity to account for cell-cycle variations and specifically to distinguish between cells in P and Q to identify and interpret intrinsic functional differences associated with acquired resistance.

### Drug resistance associates with distinct patterns of activity changes between proliferation and quiescence

As most clones and gene set activity showed the strongest differences between P and Q phases, we next summarized the observations across these two phases (Figure 4D, and Figure S7). The median activity level of cluster B, C and D was the lowest, irrespective of the phase. In contrast, the median activity in cluster E and F had levels similar to cluster A and even slightly higher in the Q phase. Across all gene sets activities, we observed three correlation patterns relative to cluster A untreated cells (Figure 4E): 1) a *P-like* pattern observed in P phase of cluster E and F, highly correlated to P phase of cluster A, 2) a *Q-like* pattern observed in Q phases of cluster E and F, correlated to Q phase of cluster A, 3) an *anti-P* pattern observed in both P and Q phases of cluster B,C and D, with activity levels with a strong negative correlations in comparison to P phase of cluster A.

The cells following the anti-P pattern all belonged to cluster B, C and D, the clusters with the most repressed activity in both P and Q phases. The gene sets that are the most active in the proliferating cells from cluster A are also the most repressed in these cells, irrespective of the cell cycle phase, suggesting the cells adopt an expression state the most functionally distant from untreated proliferative cells and more closely resembling untreated quiescent cells. The resistant phenotype in these cells could therefore be due to the maintenance of a quiescence-like state, even in cells that are proliferating. In contrast, cells from cluster E and F in P and Q phases followed the P-like and Q-like patterns respectively, suggesting they resemble more closely cluster A cells. The level of correlation with the Q phase is however weaker, suggesting the difference may contribute to the resistance. Importantly, the patterns observed are unlikely due to genetic changes or associated with resistant steps as one clone switched from Anti-P (R14 step 5) to P-like and Q-like (R14 step 15) during the course of the treatment. Furthermore, the patterns are independent from differences in proliferation rate, or fraction of cells in Q, as cells from slow proliferation (cluster B) or faster proliferation (cluster D) are both following the anti-P pattern.

The activity of some gene sets did not follow the patterns described above and their analysis may help determine which process are essential or dispensable to the resistant phenotype in specific clusters (Figure S7). Increased IFNα response signaling in both P and Q phases of cluster C suggests that this process has been activated for clone R18 between step 5 and step15 of the treatment. Cells from cluster D showed induction of Wnt-signaling in both P and Q phases and UV response in P phase to levels similar to untreated cells, suggesting these processes may not contribute to the resistant phenotype as their activity was restored during subsequent treatment of that clone. Compared to Q phase untreated cells, Q phase cells in both cluster E and F have more active inflammatory response (E and F), IFNα (E only), Wnt signaling (F only) suggesting that these processes may contribute to the resistance by altering the expression state of cells in Q phase.

## Discussion

As shown from the multiple replicates, treatment steps and gene sets analyzed, the acquisition of resistance to CBDCA is a highly heterogeneous process with the repression of proliferation, and transition to a quiescent state as a common underlying factor. As such, the variation of activity of biological process along the cell cycle, and in particular between proliferation and quiescence can confound the functional annotation of the resistant phenotype. In order to determine whether the changes in expression of specific processes mediate the resistant phenotype, we used single-cell expression profiling to virtually synchronize cells and compare the activity of biological processes along the cell cycle. This analysis revealed that processes may be altered differently between P and Q phases during the acquisition of resistance and that different clones and treatment steps may follow different patterns.

In contrast to previous studies, the use of isogenic, single-cell derived cell lines allowed the generation and comparison of multiple matching resistant and sensitive samples. This rigorous experimental design, applied to the CAOV3 cell line, which unambiguously originated from a patient with HGSOC^27^, substantially reduced experimental noise. Most experimental studies of resistance choose a continuous exposure to the drug, selecting for cells in a drug persister state which expand to a drug-tolerant persister state. In contrast, in our study we chose to mimic chemotherapy cycles, giving an opportunity for the cells to recover between treatment steps through multiple cycles. Carrying out such multi-clone, multi-step experiment offers significant challenges in the laboratory, since clones rapidly acquire variable proliferation and treatment recovery dynamics. While the source of this diversity is not completely understood, it is likely rooted in the differences in Lamarckian induction of adaptive response observed in other systems^28^.

Interestingly, sequential treatment cycles such as the one used in our study may allow multiple rounds of adaptation to two different types of transitions to occur: from drug-free to treatment (drug on) and reciprocally (drug-off), providing, therefore, multiple chances for such adaptation – and associated heterogeneity – to occur. There is a strong association between cell cycle transition and drug exposure transition, as cells exposed to the drug will activate G1 checkpoint and may enter into quiescence, until repair can be completed and drug removed. Reciprocally, drug removal eventually leads to re-entry in cell cycle. As a consequence processes active in P phase are more likely to impact the drug-on transition whereas processes active in Q phase would be more likely to impact the drug-off transition. Specifically, processes accelerating P to Q transition or slowing Q to P transition are both likely to increase resistance by keeping cells in a low proliferative state. Such a hysteresis resistance model (Figure S8) is compatible with the following observations: processes in cells from cluster B had activity levels similar to, or lower than, untreated cells in Q phase, suggesting that cells in P phase are primed for a fast P to Q transition in this earlier treatment step. Alternatively, processes in cells from cluster E and F in Q phase have higher activity levels than untreated cells in Q phase suggesting that these processes slow down Q to P transition in support of resistance.

Cellular hysteresis models similar to the one discussed here, have been used previously to describe antibiotic resistance in bacteria^29^ or transition between epithelial and mesenchymal states in cancer cells^30^, but are not commonly used to model cancer drug resistance. The model accounts for the treatment memory effect that has been observed^31^, proposing that transition between states is not symmetric and can be mediated by distinct processes. Single-cell RNA-sequencing (scRNA-seq) and virtual synchronization allowed us, for the first time, to independently study cells undergoing drug exposure in each state thereby offering an observation window on the processes already primed before a transition occurs. However, the treatment time course (step 5 and step 15) and observation window (post recovery) used in our experiments lacks sufficient resolution to fully validate the model on individual treatment cycles and to precisely follow how the activity of different processes changes as a function of drug concentration, exposure or recovery duration, or number of cycles. The collection of multiple time points, within hours before and after each transition is likely to provide much clearer information. Importantly, the use of single-cell assays such as scRNA-seq or RNA-FISH is critical to capture events shortly after treatment where only few cells remain and to alleviate the need to expand them and introduce undesirable variation.

Based on our observations, it is unlikely that a single signaling or Hallmark process exclusively contributes to the acquisition of CBDCA resistance. The bulk analysis identified IFNα response signaling as an induced process shared by multiple clones. Activation of interferon signaling is triggered by the DNA damage response (DDR) ^32^ and was previously observed in response to genotoxic stress. Expression of *IRF1*, a main effector of interferon signaling, is induced by cisplatin and may limit this drug’s effectiveness ^21^. The process could be successfully blocked by pre-exposure to ruxolitinib which restored sensitivity. Hence, consistent with the findings of the genetic profiling, this observation suggested that resistance is unlikely to be inherited. The virtual synchronization showed little variation in IFNα response signaling along the cell cycle, which suggests that despite a strong shift of resistant clones into quiescence, the bulk analysis may have measured intrinsic changes independent of cell cycle. Interestingly, the level of induction of IFNα in both P and Q phase was correlated with the response to ruxolitinib with R14 and R18 clones being the most sensitive. IFNα response was, however, not induced in cluster B corresponding to step 5 treatment, perhaps suggesting that the sensitizing effect of ruxolitinb, although visible in untreated cells, may not apply to earlier treatment steps. Furthermore, it is not clear whether the inhibition of JAK/STAT signaling would restore sensitivity by accelerating the Q to P transition or slowing down P to Q. Studies in tissue regeneration have shown that inhibition of JAK/STAT promotes stem cell expansion suggesting that inhibition of JAK/STAT could wake up quiescent cells^33^. Such considerations on the direction of the effect are important for the design of combination therapy to prevent or reverse resistance and more precisely determine the treatment schedule. One can envision combination of platinum drugs with treatments slowing down P to Q transition to prevent resistance development. Reciprocally, treatment accelerating Q to P transition, targeting processes active in Q phase, would increase the benefit of drug holidays and accelerate the re-sensitization of the tumor.

Beyond IFNα response signaling, it is possible that many other processes are impacting the dynamics of the transition at every treatment cycle and such redundancy may explain the heterogeneity observed between clones or after multiple cycles. However, cellular processes are constrained by the underlying regulatory circuitry and strongly inter-dependent. Multiple research efforts are underway to map these dependencies and reduce the complexity to fewer comprehensible dimensions^34,35^. The cellular hysteresis model of transition however adds a dynamic dimension ignored so far. The addition of single-cell assays and virtual synchronization to systematic, large scale assays will likely help generalize the phenomenon to multiple stimuli and understand which processes are acting in concert along each transition direction. It is likely that some of them may be key regulators of one direction without affecting the other. Such precise mapping would be important to determine which processes, or combination of processes, can be targeted to more effectively prevent the acquisition of resistance or more rapidly and durably re-sensitize cells. Similarly, the analysis of the epigenetic changes associated with the transitions are likely to capture underlying regulatory mechanisms supporting the hysteretic memory of change in growth conditions. Subsequent analysis at single-cell resolution would be needed to distinguish regulatory elements primed or poised for each transition direction.

Importantly, while our demonstration relies on in vitro observations, these processes are at play *in vivo*, in patients, where multiple stimuli from the micro-environmental niche, in addition to the treatment itself, may impact the cells’ decision to enter or exit a proliferative state and it is likely that the hysteresis paradigm applies to any transition between states, which in oncology, often boils down to proliferation, quiescence or cell-death.

## Materials and Methods

### Generation of CBDCA resistant clones

CAOV3 cells and all sublines were grown in RPMI 1640 containing 5% fetal bovine serum and 1X penicillin/streptomycin. The clonally derived cell lines were always plated at 40,000 cells per well in 6 well plates and allowed to attach overnight before adding the drug. A selection cycle consisted of exposure to the drug for 7 days following which cells were allowed to recover in drug-free medium for ∼2 weeks until they resumed growth and reached confluence. CBDCA sensitivity was determined from concentration-survival curves using ≥5 concentrations; viability was determined with the Cell Counting Kit 8 (Dojindo Molecular Technologies, Rockville, MD) or Crystal Violet reagent after 96 h of drug exposure.

### 2D and 3D Growth Assays

CAOV3 cells were seeded at 200 cells/well in a 6-well plate in replicates of 3 or 6 and allowed to form colonies for 9 days after which they were stained with Crystal Violet. Colonies were counted microscopically. The capacity to grow in 3 dimensions was tested by seeding 20,000 cells/well in ultra-low attachment 6 well plates (Corning Ref 3471) in stem cell medium (1:1 DMEM:F12 plus L-glutamine, 15 mM HEPES, 100 U/mL penicillin, 100 μg/mL streptomycin, 1% knockout serum replacement, 0.4% bovine serum albumin, and 0.1% insulin-transferrin-selenium (Corning, Corning, NY). The stem cell medium was further supplemented with human recombinant epidermal growth factor (20 ng/mL) and human recombinant basic fibroblast growth factor (10 ng/mL). The medium in each well was refreshed every 3 days by adding 500 µL/well of fresh stem cell media supplemented with the growth factors. Spheres were counted under a microscope and subclassified as either tight spheres or organoids after 7 and 14 days of culture.

### Ruxolitinib Treatment

#### Validation of JAK inhibition

The parental clones (6 wells seeded at 10^5^ cells/well) were pre-treated with 5µM of Ruxolitinib (50 mM in DMSO – LC laboratories) or vehicle. The cells were then treated with 1000 units of human Interferon-β for 24 hours. RNA was then isolated using TRIzol (ThermoFisher), quantified using Nanodrop and 1 µg was converted to cDNA (High Capacity cDNA Reverse Transcription Kit - Applied Biosystems). The quantitative real-time PCR amplification -95°C (10 min) and 34 cycles of 95°C (1 min), 60°C (30s), 72°C (1 min) – final: 72°C (5min) - was carried out in duplicate using the following pairs of primers: ISG15_FWD (GAGAGGCAGCGAACTCATCT), ISG15_REV (CTTCAGCTCTGACACCGACA) and 18S_FWD (CGCCGCTAGAGGTGAAATTCT) and 18S_REV (CGAACCTCCGACTTTCGTTCT). Expression data were normalized to the geometric mean of housekeeping gene 18s to control the variability in expression levels and were analyzed using the 2-ΔΔCT method.

#### Dose-Response Assay

The growth rate of each clone was first established by seeding in 500, 1000, or 2000 cell per well and monitoring for 96 hrs (3 wells per time point, per sample). The density allowing exponential growth at 96 hr was chosen to seed the cells in triplicate (96-well plates): Parent and R06: 2000 cells/well, R14: 2500 cells/well, R16: 3000 cells/well, and R18: 3000 cells/well. The cells were pre-treated with 5 μM Ruxolitinib or vehicle. Two additional wells were seeded in parallel to correct for growth rate differences (with and without Ruxolitinib) at the time of CBDCA treatment (time 0). The remaining wells were treated at time 0 with increasing CBDCA dose for 96 hours, following which the cells were detached using 0.25% Tryspin-EDTA, diluted 1:2 in trypan blue and viable cells counted using a Hemocytometer. The Growth Rate and Growth-Rate corrected half-maximal inhibitory concentration (GR50) were calculated following Hafner et al.^36^ and the *GRmetrics* bioconductor package.

### Mass Cytometry

Cells were incubated with 15 µM CBDCA for 1 h at 37°C, washed and then exposed to a 1:500 dilution of Cell-ID Intercalator ^103^Rh for 15 min at 37°C to mark the dead cells in the population. Analysis of ∼2 ×10^5^ cells from each sample was carried out on a Fluidigm Helios mass cytometer using EQ Four Element Calibration Beads for normalization. The results are presented in Table S3.

### Exome Sequencing and Analysis

The sequencing libraries were prepared and captured using SureSelect Human All Exon V4 kit (Agilent Technologies) following the manufacturer’s instructions. The sequencing was performed using the Illumina HiSeq 2000 system, generating 100 bp paired-end reads. All raw 100 bp paired-end reads were aligned to the human genome reference sequence (hg19) using BWA ^37^ and further jointly realigned around indels sites using GATK’s ^38^ IndelRealigner. Duplicate reads were removed using Picard Tools MarkDuplicates ^39^. Table S4 presents the summary statistics of the sequencing. The variants were called using Freebayes ^40^ and filtered for high quality (QUAL/AO>10). We annotated the variants with ANNOVAR ^41^, removed non-coding and synonymous variants, variants in dbSNP147 or shared between all samples, leading to a total of 93 variants across all 8 samples (Table S2). The copy number changes were called independently on each chromosome using CODEX ^42^ with default settings (Table S1), limited to the expected exonic target from the SureSelect capture kits and expecting fractional copy number from aneuploidy. Segments smaller than 100kb, supported by less than 3 exons, or with copy number between 1.5 and 2.5 were excluded.

### RNA Sequencing and Analysis

RNA was extracted using Qiagen RNAEasy and the libraries were prepared from 1 µg of RNA using TruSeq following the manufacturer instructions (shear time modified to 5 min). The libraries were sequenced on HiSeq 4000 (paired end 100 nt reads) and analyzed using BCBio-nextgen 1.0.1 ^43^ RNA-Seq default pipeline which included adapter removal with cutadapt v1.12 ^44^, read splice aware alignment with Bowtie2/Tophat suite v2.22.8 ^45,46^ for quality control, and isoform expression level estimation using sailfish 0.10.1 ^47^. The differential expression was determined using DESeq2 ^48^. We performed Gene Set Enrichment Analysis ^49^ implemented in the *liger* R package using gene sets from MSigDB ^19^.

### Single cell RNA-Sequencing

#### Data generation

We used the 10x Chromium (10x Genomics v2 reagents) to isolate ∼2000 single cells from each sample following the manufacturer’s instructions. Briefly, the cells/GEM droplets emulsion was formed using the 10x Chromium controller. The reverse transcription and template switching steps added both a cell-specific barcode and unique molecular identifier to each cDNA. The emulsion was then broken up and the GEM cleaned up. The single-strand cDNA was fragmented enzymatically and subjected to library preparation, including clean-up, end-repair, adapter ligation and enrichment PCR to add a sample-specific index. The libraries were quantified using Agilent Tape-station, and pooled for sequencing on the Illumina HiSeq 4000 for single index paired-end sequencing (28+98nt reads). The resulting sequencing reads were separated using bcl2fastq and analyzed using the Cell Ranger v2 pipeline *count*, combining reads from different sequencing runs.

#### Data analysis

The barcode/cell matrices from different samples were further aggregated using Cell Ranger *aggregate* normalizing to total number of reads (Table S5). The aggregated count matrices were processed with Seurat version 3.1.1^50^. Genes expressed in fewer than three cells were removed. Additionally, cells were removed if fewer than 200 genes or more than 3700 genes were detected, or if mitochondrial genes made up more than 10% of their transcriptome. The remaining data was log normalized and scaled using the *NormalizeData* and *ScaleData* functions in Seurat with default parameter settings. To identify clusters, Seurat *FindVariableGenes* and *RunPCA* functions were run with default parameters and followed by *FindNeighbors* with the top 50 principal components and a resolution of 0.2 to identify the cell clusters. To assign cells to cell cycle phases, the Seurat *CellCycleScoring* function was modified to accept more than the 2 cell cycle gene sets. Using this function, each cell was scored for six cell cycle related gene sets derived from Xue *et al*. (G0.1 gene set excluded)^51^ and was assigned the cell-cycle phase corresponding to the gene set with the maximum score.

#### Pseudo-time Gene Set Enrichment analysis

The expression of 603 genes included in the six cell cycle gene sets previously used^51^ was used to organize cells along a linear pseudo-time trajectory using SCORPIUS version 1.0.2 ^52^, following the default workflow from the tutorial^53^. Cells from each cluster were grouped in smoothing pseudo-time windows (interval=0.2, increment=0.01 as a fraction of the pseudo-time range). The gene expression of all cells in a window was summarized into a virtual cell gene expression using a method derived from Baran et al^54^. In brief, the geometric mean of each gene’s expression was calculated, divided by the median across all windows and clusters, and log transformed. The resulting gene expression profile of each virtual cell was then used to calculate the enrichment score of each gene set in the MSigDB’s Hallmark collection^17^ using single sample GSEA (ssGSEA)^55^ as implemented in the GenePattern^56^ module.

## Acknowledgements

We thank Drs. Pablo Tamayo and Xin Lian Zhang for helpful discussions. We thank Mr. Gerald Manorek for technical assistance as well as Dr. Lyn Hedrick and Erik Ehinger for assistance with the Mass Cytometry assays. The work was supported by grants from the National Cancer Institute (R21CA177519, U24CA220341, U24CA194107, U24CA248457), from the Department of Defense (OC140179), grants from The Hartwell Foundation and The Immunotherapy Foundation, and an award from Pedal the Cause and a grant from the UC San Diego Academic Senate. ATW was supported by an NLM T-15 training grant (T15LM011271) and a Training Fellowship from UC San Diego Cancer Cell Map Initiative (U54CA209891).

## Data availability

The data generated are available at the NCBI under bioproject PRJNA419934

## Author contributions

DC, SS, C-YT, HV conducted the experiments, DC, OH, ATW, JPM analyzed the data, ATW, DC, JDB, SBH, JPM and OH interpreted the data and SBH, OH, wrote the manuscript.

## References

1. Ozols, R. F. et al . Phase III trial of carboplatin and paclitaxel compared with cisplatin and paclitaxel in patients with optimally resected stage III ovarian cancer: a Gynecologic Oncology Group study. J. Clin. Oncol. 21, 3194–3200 (2003).

2. Siegel, R., Naishadham, D. & Jemal, A. Cancer statistics, 2013. CA. Cancer J. Clin. 63, 11–30 (2013).

3. Andrews, P. A., Jones, J. A., Varki, N. M. & Howell, S. B. Rapid emergence of acquired cis-diamminedichloroplatinum(II) resistance in an in vivo model of human ovarian carcinoma. Cancer Commun. 2, 93–100 (1990).

4. Galluzzi, L. et al. Molecular mechanisms of cisplatin resistance. Oncogene 31, 1869–1883 (2012).

5. Sakai, W. et al. Secondary mutations as a mechanism of cisplatin resistance in BRCA2-mutated cancers. Nature 451, 1116–1120 (2008).

6. Brown, R., Curry, E., Magnani, L., Wilhelm-Benartzi, C. S. & Borley, J. Poised epigenetic states and acquired drug resistance in cancer. Nat. Rev. Cancer 14, 747–753 (2014).

7. Sharma, S. V et al. A chromatin-mediated reversible drug-tolerant state in cancer cell subpopulations. Cell 141, 69–80 (2010).

8. Gascoigne, K. E. & Taylor, S. S. Cancer Cells Display Profound Intra- and Interline Variation following Prolonged Exposure to Antimitotic Drugs. Cancer Cell 14, 111–122 (2008).

9. Cohen, A. A. et al. Dynamic Proteomics of Individual Cancer Cells in Response to a Drug. Science (80-.). 322, 1511–1516 (2008).

10. Shaffer, S. M. et al. Rare cell variability and drug-induced reprogramming as a mode of cancer drug resistance. Nature 546, 431–435 (2017).

11. Kondoh, E. et al. Targeting slow-proliferating ovarian cancer cells. Int. J. Cancer 126, NA-NA (2009).

12. Shah, M. A. & Schwartz, G. K. Cell cycle-mediated drug resistance: an emerging concept in cancer therapy. Clin. Cancer Res. 7, 2168–2181 (2001).

13. Abada, P. & Howell, S. B. Regulation of Cisplatin cytotoxicity by cu influx transporters. Met. Based. Drugs 2010, 317581 (2010).

14. Kircher, M. et al. A general framework for estimating the relative pathogenicity of human genetic variants. Nat. Genet. 46, 310–5 (2014).

15. Zhang, Y. et al. Identification of a conserved anti-apoptotic protein that modulates the mitochondrial apoptosis pathway. PLoS One 6, e25284 (2011).

16. Aslam, M. A. et al. Towards an understanding of C9orf82 protein/CAAP1 function. PLoS One 14, e0210526 (2019).

17. Liberzon, A. et al. The Molecular Signatures Database (MSigDB) hallmark gene set collection. Cell Syst. 1, 417–425 (2015).

18. Fabregat, A. et al. The Reactome Pathway Knowledgebase. Nucleic Acids Res. (2017). doi:10.1093/nar/gkx1132

19. Liberzon, A. et al. Molecular Signatures Database (MSigDB) 3.0. Bioinformatics (2011). doi:10.1093/bioinformatics/btr260

20. Brzostek-Racine, S., Gordon, C., Van Scoy, S. & Reich, N. C. The DNA Damage Response Induces IFN. J. Immunol. 187, 5336–5345 (2011).

21. Pavan, S., Olivero, M., Corà, D. & Di Renzo, M. F. IRF-1 expression is induced by cisplatin in ovarian cancer cells and limits drug effectiveness. Eur. J. Cancer 49, 964–73 (2013).

22. Yang, A. D. et al. Chronic Oxaliplatin Resistance Induces Epithelial-to-Mesenchymal Transition in Colorectal Cancer Cell Lines. Clin. Cancer Res. 12, 4147 LP–4153 (2006).

23. Marchion, D. C. et al. BAD phosphorylation determines ovarian cancer chemosensitivity and patient survival. Clin. Cancer Res. 17, 6356–6366 (2011).

24. Cui, W., Yazlovitskaya, E. M., Mayo, M. S., Pelling, J. C. & Persons, D. L. Cisplatin-induced response of c-jun N-terminal kinase 1 and extracellular signal--regulated protein kinases 1 and 2 in a series of cisplatin-resistant ovarian carcinoma cell lines. Mol. Carcinog. 29, 219–228 (2000).

25. Persons, D. L., Yazlovitskaya, E. M., Cui, W. & Pelling, J. C. Cisplatin-induced activation of mitogen-activated protein kinases in ovarian carcinoma cells: inhibition of extracellular signal-regulated kinase activity increases sensitivity to cisplatin. Clin. Cancer Res. 5, 1007–1014 (1999).

26. Pénzváltó, Z. et al. MEK1 is associated with carboplatin resistance and is a prognostic biomarker in epithelial ovarian cancer. BMC Cancer 14, 837 (2014).

27. Beaufort, C. M. et al. Ovarian cancer cell line panel (OCCP): clinical importance of in vitro morphological subtypes. PLoS One 9, e103988 (2014).

28. Su, Y. et al. Single-cell analysis resolves the cell state transition and signaling dynamics associated with melanoma drug-induced resistance. Proc. Natl. Acad. Sci. 114, 13679 LP–13684 (2017).

29. Roemhild, R. et al. Cellular hysteresis as a principle to maximize the efficacy of antibiotic therapy. Proc. Natl. Acad. Sci. 115, 9767 LP–9772 (2018).

30. Celià-Terrassa, T. et al. Hysteresis control of epithelial-mesenchymal transition dynamics conveys a distinct program with enhanced metastatic ability. Nat. Commun. 9, 5005 (2018).

31. Bell, C. C. & Gilan, O. Principles and mechanisms of non-genetic resistance in cancer. Br. J. Cancer 122, 465–472 (2020).

32. Nakad, R. & Schumacher, B. DNA Damage Response and Immune Defense: Links and Mechanisms. Front. Genet. 7, 147 (2016).

33. Price, F. D. et al. Inhibition of JAK-STAT signaling stimulates adult satellite cell function. Nat. Med. 20, 1174–1181 (2014).

34. Tsherniak, A. et al. Defining a Cancer Dependency Map. Cell 170, 564–576.e16 (2017).

35. Kim, J. W. et al. Decomposing Oncogenic Transcriptional Signatures to Generate Maps of Divergent Cellular States. Cell Syst. 5, 105–118.e9 (2017).

36. Hafner, M., Niepel, M., Chung, M. & Sorger, P. K. Growth rate inhibition metrics correct for confounders in measuring sensitivity to cancer drugs. Nat. Methods 13, 521–527 (2016).

37. Li, H. & Durbin, R. Fast and accurate short read alignment with Burrows-Wheeler transform. Bioinformatics 25, 1754–1760 (2009).

38. McKenna, A. et al. The Genome Analysis Toolkit: a MapReduce framework for analyzing next-generation DNA sequencing data. Genome Res. 20, 1297–303 (2010).

39. Picard. Available at: http://sourceforge.net/projects/picard/.

40. Garrison, E. & Marth, G. Haplotype-based variant detection from short-read sequencing. 9 (2012).

41. Wang, K., Li, M. & Hakonarson, H. ANNOVAR: functional annotation of genetic variants from high-throughput sequencing data. Nucleic Acids Res. 38, e164 (2010).

42. Jiang, Y., Oldridge, D. A., Diskin, S. J. & Zhang, N. R. CODEX: a normalization and copy number variation detection method for whole exome sequencing. Nucleic Acids Res. 43, e39 (2015).

43. bcbio-nextgen. Available at: https://github.com/chapmanb/bcbio-nextgen.

44. Martin, M. Cutadapt removes adapter sequences from high-throughput sequencing reads. EMBnet.journal; Vol 17, No 1 Next Gener. Seq. Data Anal. (2011).

45. Langmead, B. & Salzberg, S. L. Fast gapped-read alignment with Bowtie 2. Nat Meth advance on, (2012).

46. Kim, D. et al. TopHat2: accurate alignment of transcriptomes in the presence of insertions, deletions and gene fusions. Genome Biol. 14, R36 (2013).

47. Patro, R., Mount, S. M. & Kingsford, C. Sailfish enables alignment-free isoform quantification from RNA-seq reads using lightweight algorithms. Nat. Biotechnol. 32, 462–464 (2014).

48. Love, M. I., Huber, W. & Anders, S. Moderated estimation of fold change and dispersion for RNA-seq data with DESeq2. Genome Biol. 15, 550 (2014).

49. Subramanian, A. et al. Gene set enrichment analysis: A knowledge-based approach for interpreting genome-wide expression profiles. Proc. Natl. Acad. Sci. 102, 15545–15550 (2005).

50. Stuart, T. et al. Comprehensive integration of single cell data. bioRxiv 460147 (2018). doi:10.1101/460147

51. Xue, J. Y. et al. Rapid non-uniform adaptation to conformation-specific KRAS(G12C) inhibition. Nature 577, 421–425 (2020).

52. Cannoodt, R. et al. SCORPIUS improves trajectory inference and identifies novel modules in dendritic cell development. bioRxiv 79509 (2016). doi:10.1101/079509

53. Cannoodt, R. SCORPIUS. Available at: https://github.com/rcannood/SCORPIUS.

54. Baran, Y. et al. MetaCell: analysis of single-cell RNA-seq data using K-nn graph partitions. Genome Biol. 20, 206 (2019).

55. Barbie, D. A. et al. Systematic RNA interference reveals that oncogenic KRAS-driven cancers require TBK1. Nature 462, 108–112 (2009).

56. Reich, M. et al. GenePattern 2.0. Nature genetics 38, 500–501 (2006).

