## Supplemental Figures for "Single-cell characterization of step-wise acquisition of carboplatin resistance in ovarian cancer"

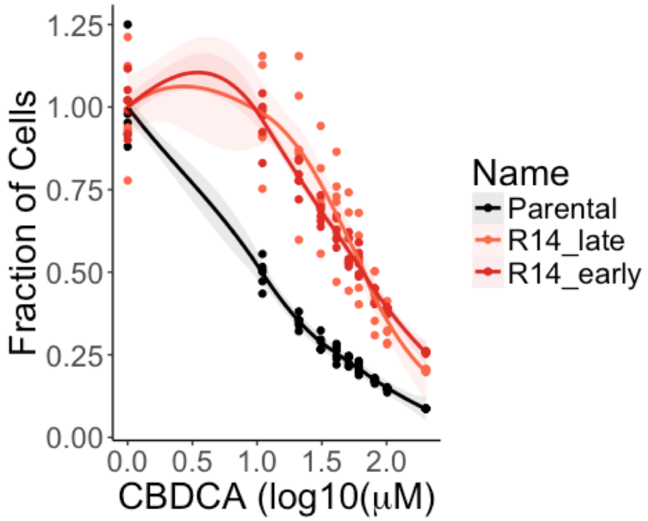


**Figure S1:** Dose Response comparing the drug sensitivity of R14 clones after 36 (R14_early) and 99 (R14_late) doublings after step 15 selection.

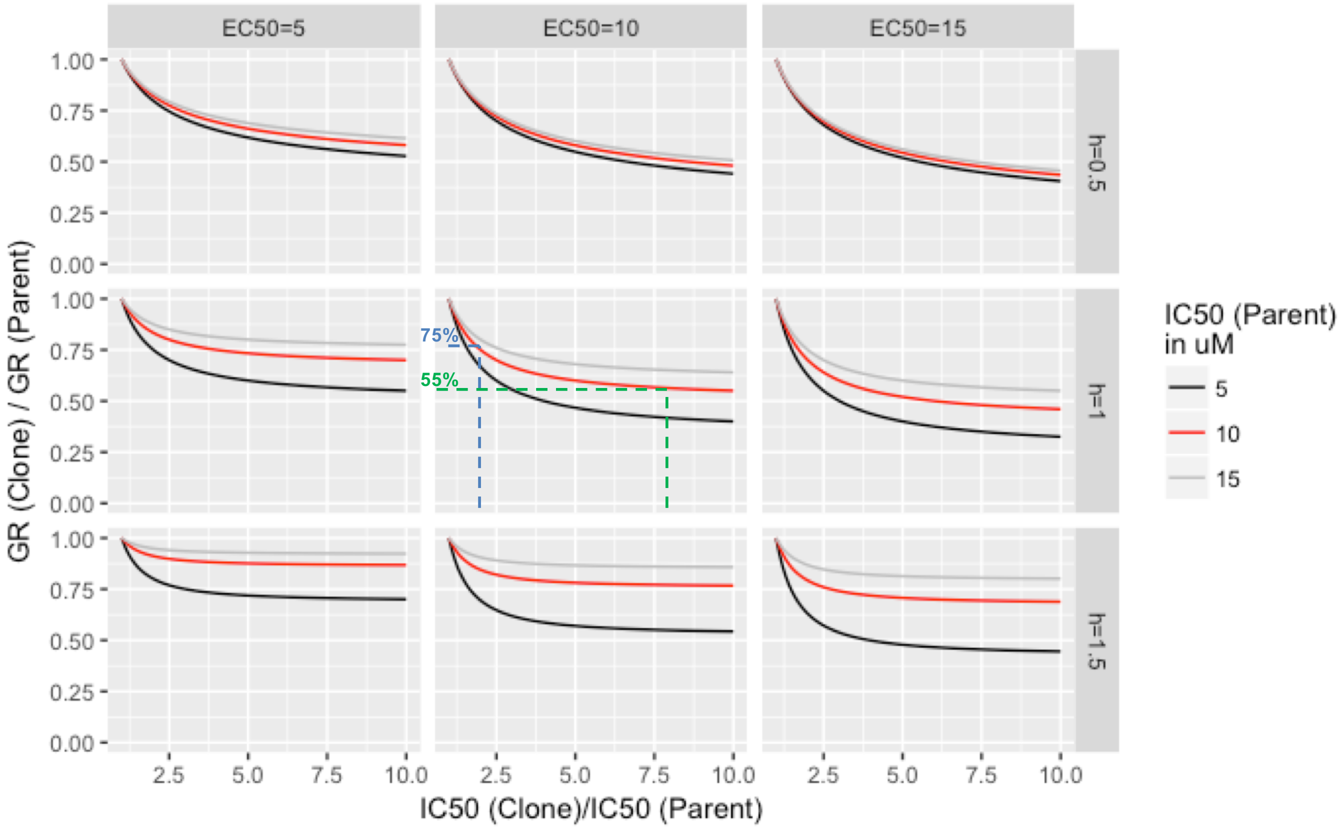


**Figure S2: Simulated impact of growth rate on resistance.** The reduction in growth rate (y axis) is associated with an increase in IC50 (x-axis). The simulations were performed according to Hafner et al (formula IC(c,t) located at http://www.grcalculator.org/grtutorial/). The IC50 of the parental clone (colored lines), hill coefficient (grid rows) and EC50 (grid columns) were tested around the experimental ones observed (IC50=10 μM, h=1, EC50=10 μM, middle panel of the grid, red line). The level of resistance observed in step 5 (blue lines) and step 15 (green lines) is indicated.

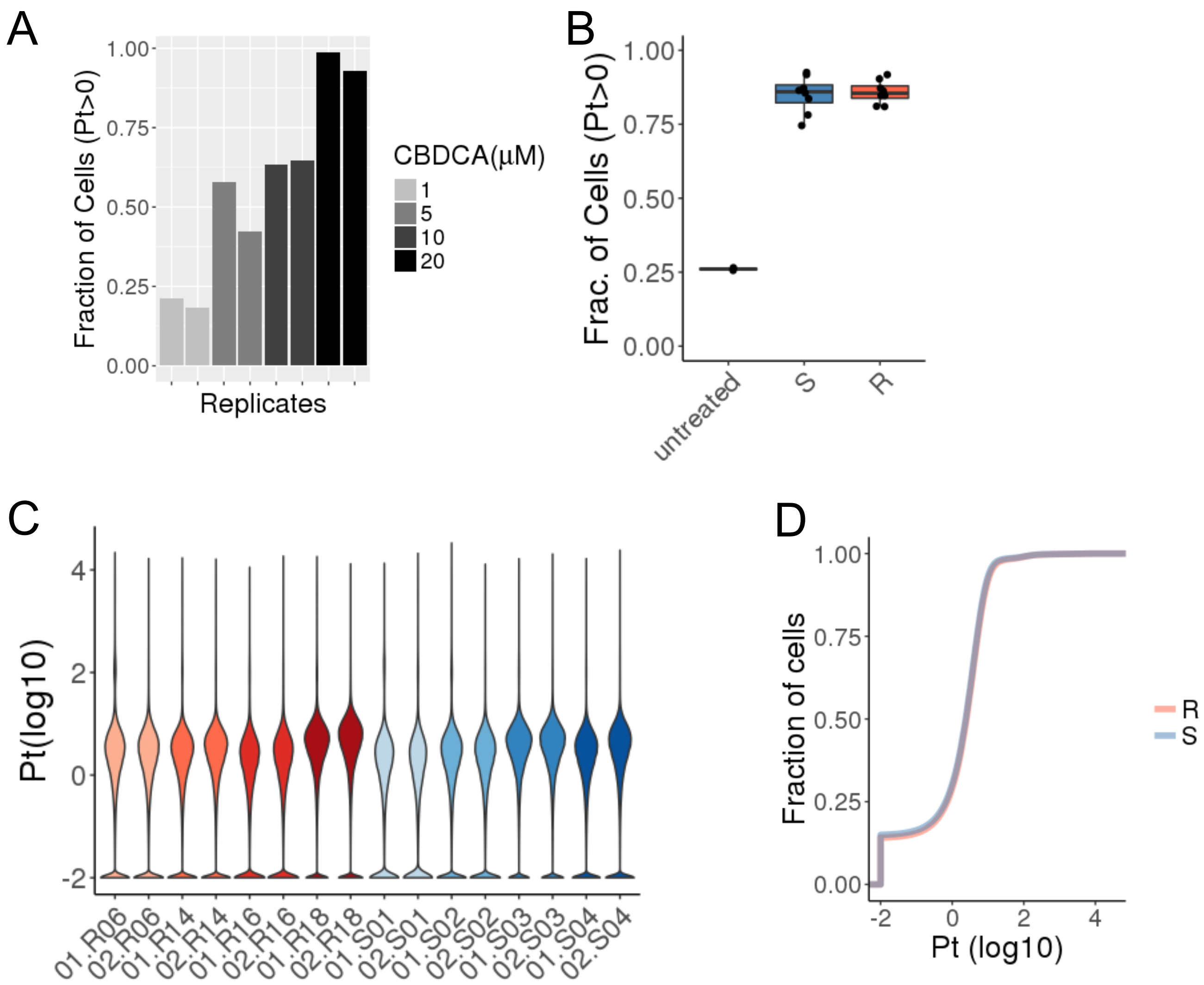


**Figure S3: Platinum uptake measured by mass cytometry. (A)** The fraction of cells form the parental clone with detectable Pt content was measured after 1 h treatment with increasing dose of carboplatin (x axis, grey color gradient). **(B)** Fraction of cells with detectable levels of Pt. Each clone (S and step 15 R clones) and replicate are represented. **(C)** Violin plot showing the distribution of Pt content across cells from all clones and replicates. **(D)** Cumulative distribution of Pt content between cells from S clones and step 5 R clones and replicates. The Pt negative cells were assigned at 0.01 Pt content for graphical representation (C and D).

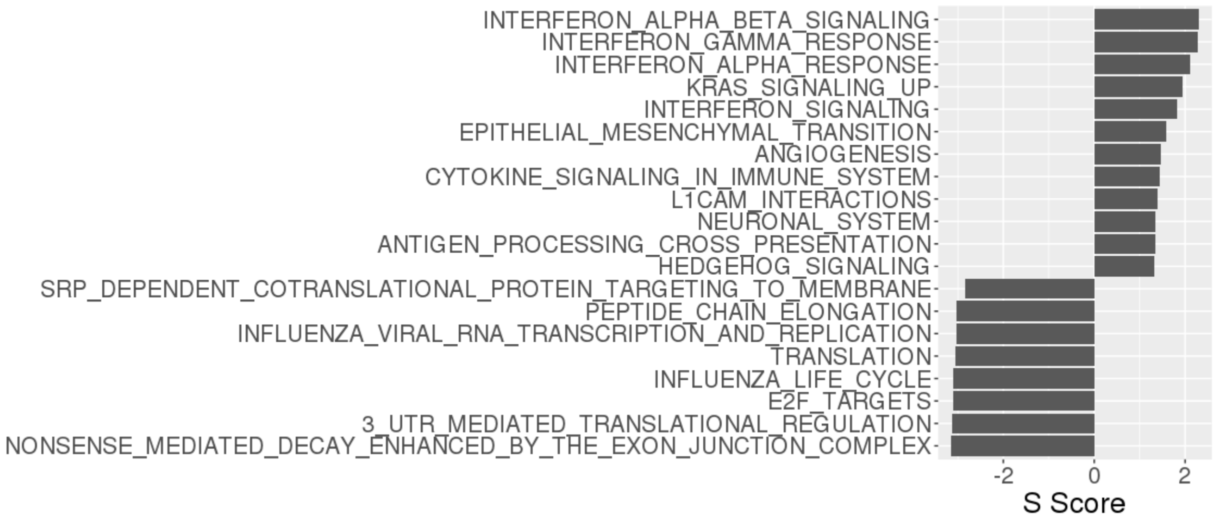


**Figure S4: Most affected pathways in the resistant clones.** Most significantly up or down-regulated processes in CBDCA resistant clones after step 15 selection (q.value<0.05). All Hallmark (N=50) and Reactome (N=674) gene sets from MSigDB were included in the analysis.

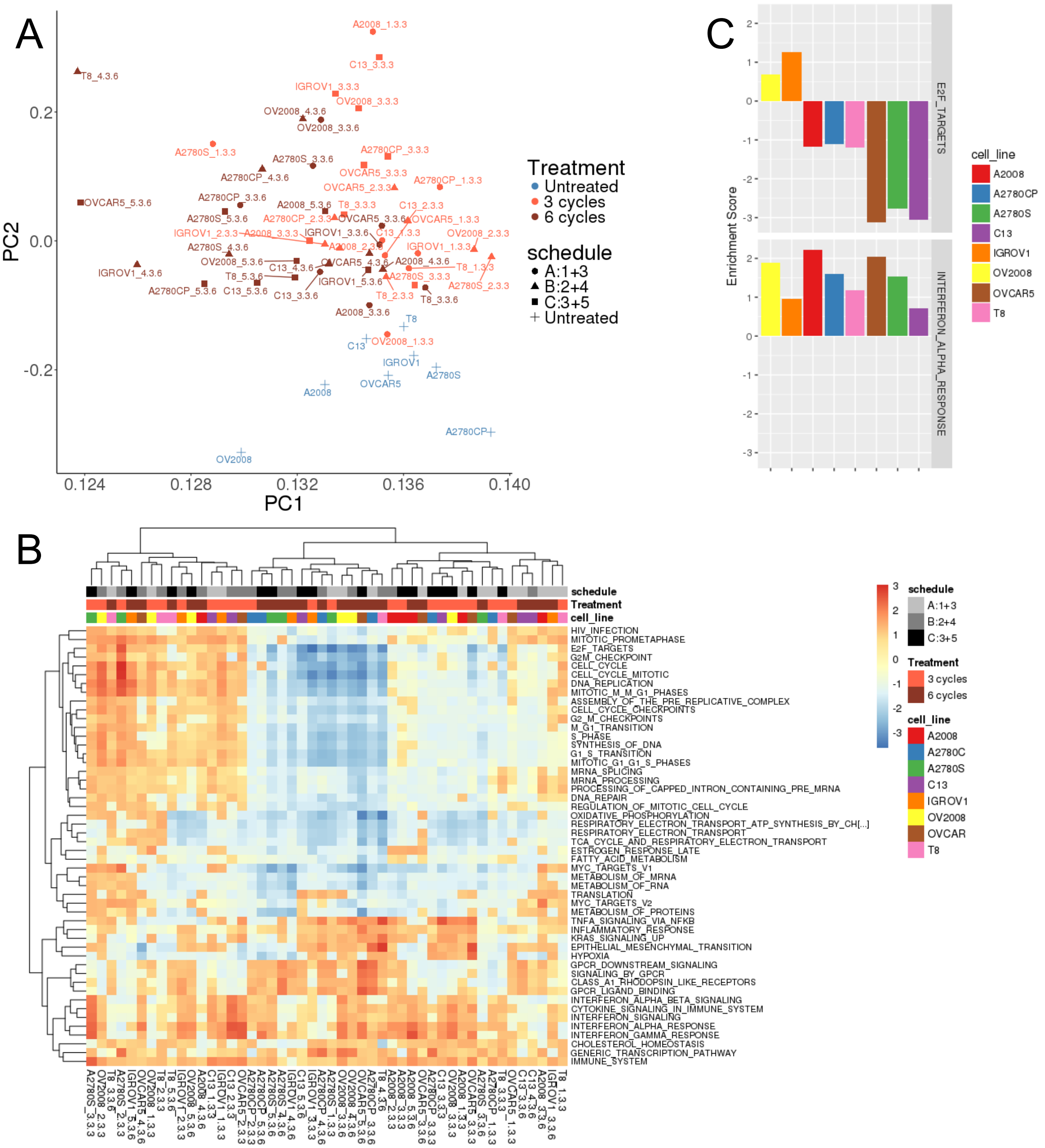


**Figure S5: Signatures of acquired cisplatin resistance in 8 ovarian cancer cell lines. (A)** Principal component analysis of 56 samples from 8 cell lines. Resistant lines were derived from each cell line using 3 treatment schedules of 6 cycles each, increasing cisplatin concentration after 3 cycles. Cells were analyzed after 3 and 6 treatment cycles. Cell line specific variance was corrected for batch effect. See Methods and Marchion et al for details (**ref**). **(B)** Fold change in gene set expression levels compared to all untreated samples. In each treated sample and the set of 8 untreated samples was used to calculate the enrichment score of all Hallmarks and Reactome gene sets from MSigDB. Significant gene sets more than 4 fold induced or repressed in at least one sample are represented. The color scale represents the enrichment score (log scale). **(C)** Enrichment score for two representative gene sets for samples after 6 cycles of treatment schedule C (3 μM then 5 μM)

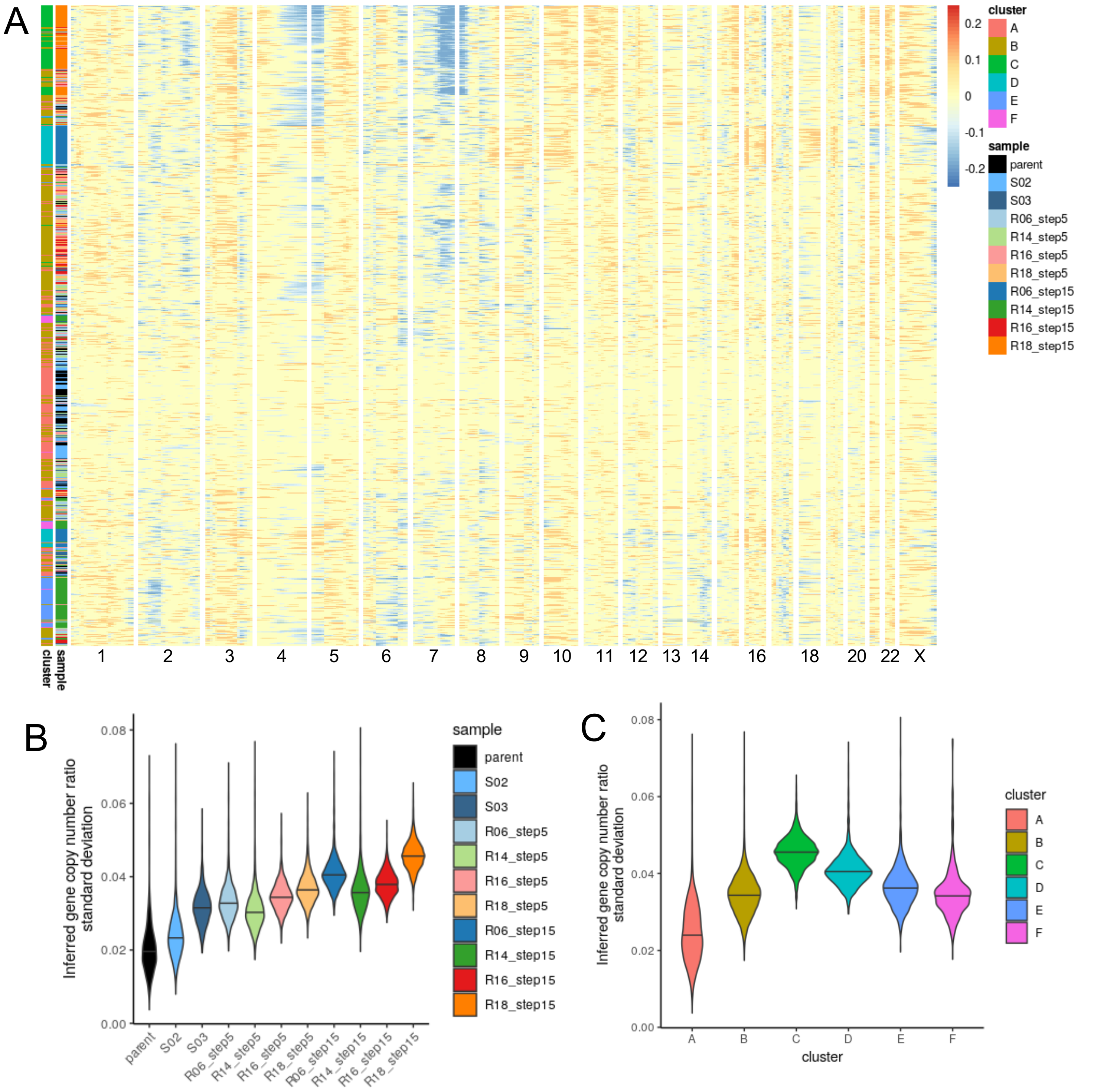


**Figure S6: Chromosome Copy Number Profile inferred from scRNA-seq expression. (A)** clustered heatmap of the copy number ratio observed in individual cells (rows) annotated based on their sample and cluster assignment. Genomic interval bins (columns – 10 Mbp windows) are group by chromosome (grouped columns). Values represent the log2 of the average copy number. The COAV3 cells were used as reference group. **(B-C)** Distribution of the standard deviation in gene copy number ratio for all cells in each sample (B) or expression-based cluster (C)

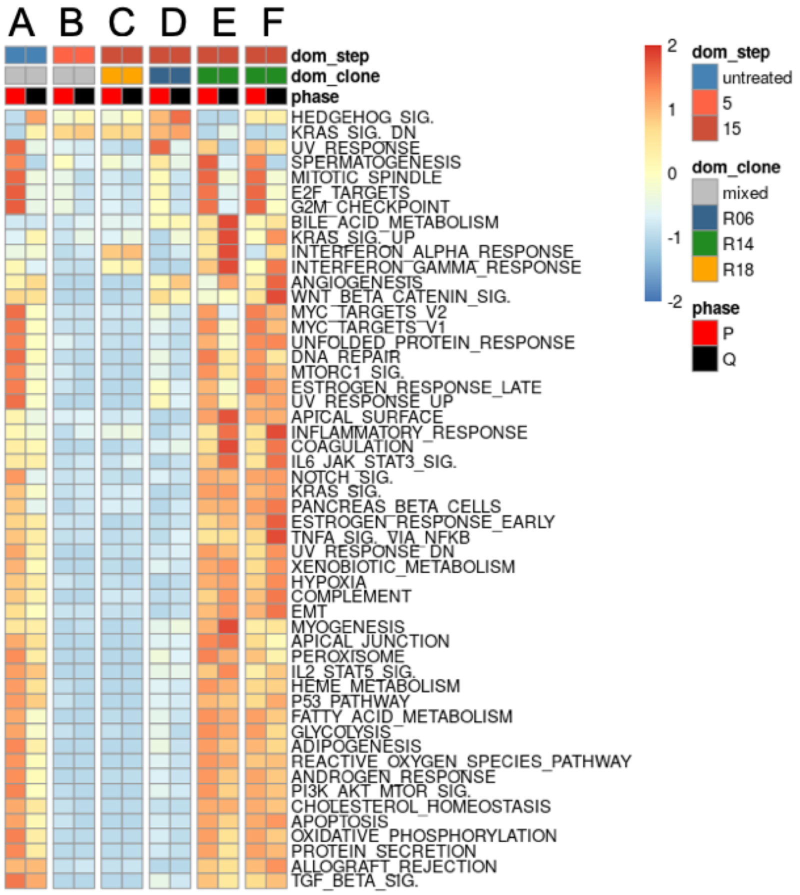


**Figure S7: Clustered heatmap** representing the median gene set enrichment scores (z-score - color-scale) for hallmark gene sets (rows) for all clusters in proliferative (P) or quiescence (Q) phases. Dominant treatment step (dom_step) and clone (dom_clone) in each cluster are indicated.

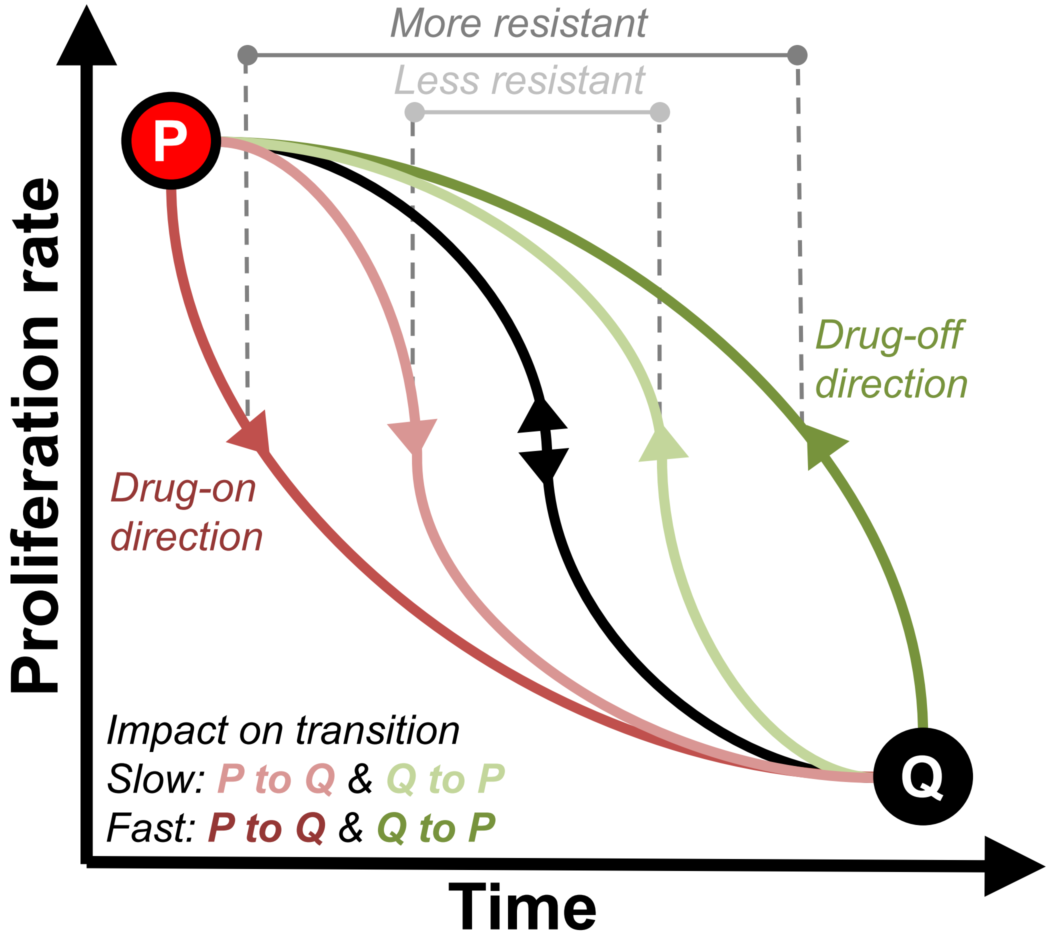


**Figure S8: A hysteresis model of resistance.** Transition between proliferation (P) and quiescence (Q) occurs at each treatment cycle following directional transitions on the hysteresis diagram: drug-on (red, downward paths) and drug-off (green upward paths) transitions. The functional state of the cells in P and Q, different after each cycle, impacts the effectiveness of the transition: light colors and low time-wise slope: slow path, dark color and steep time-wise slope: fast path.
